# Type-I interferons drive the gastrointestinal inflammatory response in a mouse model of Parkinson’s disease

**DOI:** 10.1101/2024.05.05.592614

**Authors:** Harrison Waters, Shuyan Chen, Elizabeth Vincan, Dustin J. Flanagan, Renate H.M. Schwab, Peter J. Crack, Juliet M. Taylor

## Abstract

**Background and Aims:** Parkinson’s disease (PD) is an age-related neurodegenerative disorder characterised by classical motor symptoms due to a loss of dopaminergic neurons in the substantia nigra pars compacta. The type-I interferons (IFNs) are elevated in the aging brain and we have implicated them in the neuroinflammatory response in PD. With increasing evidence of gastrointestinal (GI) dysfunction in PD patients, this study explored the contribution of the type-I IFNs to the transmission of pathology from the brain to the gut in PD.

**Methods:** Young (10-12 weeks) and aged (40-50 weeks) wildtype and IFNAR1^−/−^ mice received an intrastriatal injection of human alpha-synuclein (α-Syn) pre-formed fibrils (PFF) (8ug) with gut tissue analysed 6-months post-injection (p.i). A mouse intestinal organoid culture model was established to further characterise the α-Syn induced inflammatory response in the gut.

**Results:** An intrastriatal injection of human α-Syn PFFs was shown to initiate a type-I IFN-dependent neuroinflammatory response in the GI tract of wildtype mice at 6-months p.i. This response was attributed to an elevation in type-I IFN signalling in aged mice that was absent in the IFNAR1^−/−^ mice. Mouse intestinal organoid cultures confirmed α-Syn was taken up by the enteroendocrine cells (EECs) to induce a type-I IFN mediated pro-inflammatory response that was attenuated in IFNAR1^−/−^ cultures.

**Conclusion:** This study has confirmed the type-I IFNs modulate the α-Syn PFF induced inflammatory response within the gut potentiating pathology progression along the gut-brain axis. Early intervention of this type-I IFN response may be a potential therapeutic target to limit the progression of PD.

## Introduction

Parkinson’s disease (PD) is a chronic, progressive neurodegenerative disorder that affects approximately 6.9 million individuals worldwide^1^. Aging is considered the largest risk factor for developing PD and with an aging population, this number is expected to reach 14.2 million in 2040.^1^ The major clinical symptoms of PD are those associated with motor dysfunction such as tremor, bradykinesia, postural instability, and muscle rigidity.^2^ However, PD patients also present with non-motor symptoms including sensory dysfunction, constipation, speech issues, hypotension, sleep disorders, difficulty swallowing, cognitive changes, and mental health issues.^3^ Some of these symptoms, such as olfactory dysfunction, constipation, and sleep disorders may present years before diagnosis. Specifically, GI disturbances, such as constipation, can predate common motor symptoms by up to 20 years.^4^ The motor symptoms have been attributed to the degeneration of the dopaminergic neurons within the substantia nigra, however the precise mechanisms underlying the disease pathogenesis remain unclear.^5^ A key pathological hallmark feature within the PD brain is the presence of Lewy body’s containing aggregated α-Synuclein (α-Syn). However, the brain is not the only region of the body that has been discovered to develop α-Syn inclusions and Lewy pathology. The identification of α-Syn aggregates within the GI tract^6^ provides a direct link between the gut and brain in the progression of the disease.^7^ Specifically, colorectal biopsies of PD patients display phosphorylated α-Syn (p-α-Syn) pathology, and colonic submucosa p-α-Syn levels have been detected in early stage untreated PD patients. ^8, 9^ Moreover, the expression of phosphorylated-α-Syn in enteric neurons has been shown to correlate with acute and chronic inflammation of the intestinal wall. The density of neutrophils and mononuclear cells is elevated in patient biopsy specimens with elevated numbers of CD68-positive cells within the lamina propria.^10^ This suggests that similar to the CNS, α-Syn induces an inflammatory response in the ENS, however the mechanisms involved are yet fully understood.

Type-I interferons (IFN), a family of pleiotropic cytokines, have been shown to be upregulated in the SNpc of post-mortem PD.^11^ Furthermore, the presence of this upregulated IFN signature in neurodegenerative diseases has been linked to a ‘reactive’ microglial profile. Specifically, single-cell RNA sequencing has identified subpopulations of microglial cells coined “interferon response microglia” that are increased in Alzheimer’s disease and in the aged brain ^12, 13^. The type-I IFNs in have also been shown to play a role in the neuroinflammatory response in genetic mouse models of PD. *Prkn^−/−^*and *Pink1^−/−^* mice display a significant inflammatory phenotype after exhaustive exercise that has been shown to be completely rescued through the loss of STING (Stimulator of Interferon Genes), a downstream mediator of the type-I IFN pathway.^14^ Furthermore, the loss of dopaminergic neurons in the SNpc and subsequent motor deficits seen in *Prkn^−/−^* mice were shown to be rescued by targetting STING. In the gastrointestinal tract, the type-I IFNs have been shown to regulate intestinal barrier function with the genetic ablation of IFNAR1 leading to intestinal hyperplasia, and a lethal loss of intestinal barrier function.^15^ With the type-I IFNs driving neuroinflammation in the PD brain and also playing a crucial role in maintaining intestinal barrier function, we hypothesised that they modulate the inflammatory response in the gut in PD, thereby contributing to the disease progression along the gut-brain axis. This study confirmed that following an intra-striatal injection of a-Syn PFFs, an inflammatory response and gut dysfunction was induced that was mediated by the type-I IFNs. The modulation of this a-Syn induced inflammatory response was confirmed *in vitro* using gut organoid cultures identifying a potential new pathway to target early non-motor symptoms and disease progression in PD.

## Materials and Methods

### Animals and ethics

All experimental procedures were conducted in compliance with the guidelines of the National Health and Medical Research Council (NHMRC) of Australia for animal experimentation. All procedures using mice were approved by the Animal Ethics Committee of the University of Melbourne (Ethics #22740). Age matched male and female C57BL/6J and IFNAR1^−/−^ mice were used for primary and organoid cultures (Ethics #22740). All mice were housed in the Biomedical Sciences Animal Facility, The University of Melbourne. Mice were maintained at 22±2°C on a 12:12 hour day/night cycle and fed a standard sterile diet of mouse chow with water available ad libitum in standard specific pathogen-free micro isolator cages. All experimental work involving animals followed the ARRIVE guidelines for animal research.

### Fibrilization of α-synuclein peptide and Thioflavin-T assay

1mg synuclein (rPeptide Inc) was diluted to 4mg/ml in sterile endo-toxin free dPBS (Dulbecco’s PBS, Ca2+/Mg2+ free) and incubated at 37°C with agitation on an orbital mixer (1000rpm) for 7 days. A Thioflavin T (ThT) assay was used to determine α-synuclein aggregation using a FLEX3B station (excitation 450nM, emission 485nm).

### Intrastriatal injection of human a-synuclein pre-formed fibrils

WT (C57Bl/6J) and IFNAR1^−/−^ (young: 8-12-weeks old and aged: 40–50-weeks old, mixed male and female) mice were anesthetized via isoflurane (ISOFLURIN®) (2-5%) inhalation. Mice were transferred onto a digital stereotaxic frame and placed in a nose cone for steady isoflurane inhalation. After drilling a 1mm burr-hole in the skill, a 5µl Hamilton syringe with either dPBS (Mg^2+^/Ca^2+^ free) (Gibco™, 14200075) or 4mg/ml α-Syn PFF (rPeptide®) was used to deliver 2µl, at a rate of 0.2µl/min, into the right dorsal striatum at the following stereotaxic coordinates relative to bregma (mm): −2.0 ML, +0.5 AP, −3.0 DV (**Supp Figure 1**). Mice were given a subcutaneous injection of 0.1mg/kg Buprenorphine (120µl) (Lyppard Australia Pty Ltd) and sterile lactated Ringer’s solution (1ml) (Thermo Fischer Scientific) to facilitate recovery. Wounds were closed using 9mm Reflex clips. Post-operative mice were closely monitored with a further 0.1mg/kg Buprenorphine administered after 24 hours.

### Isolation of intestinal crypts

Intestinal crypts were isolated from eight-week-old C57BL/6J and IFNAR1^−/−^ mice using the method of Flanagan et al. (2019).^16^

### Immunohistochemical analysis of intestinal organoid cultures

WT and IFNAR1^−/−^ intestinal organoids were grown in 24-well plates, in 50μl BME-2 spheres. Organoids were fixed with ice-cold 4% w/v paraformaldehyde and gently pipetted up and down to dissociate the BME-2. The organoids were transferred to a 1.5ml Eppendorf tube and filled to 1ml with 4% w/v paraformaldehyde which was then incubated on ice for 30 minutes. The organoids were allowed to pellet via gravity and then washed with 1xPBS (3 x 10 mins). The PBS was removed and replaced with PBSDT Blocking solution (1XPBS supplemented with 0.1-1% Triton X-100, 1% DMSO, 1% BSA, and 1% Goat serum) (500μl, gentle agitation, RT, 2 hours) with the tube allowed to stand for 10 mins for the organoids to again pellet via gravity. The PBSDT was then removed and replaced with 10% v/v CAS-Block™ (008120, Invitrogen, Scoresby, VIC, Australia). The blocking agent was then removed and replaced with primary antibodies **(Supp Table 1)** diluted in PBSDT for 24 hours with gentle agitation. Following an overnight incubation, the cells were washed with PBS/0.1%BSA (4 x 10 mins), the supernatant was then removed, and the organoids incubated with secondary antibody **(Supp Table 1)** diluted in PBS/0.1%BSA (gentle agitation, RT, 2 hours). The cells were washed with PBS/0.1%BSA (4 x 10 mins) before and additional wash with 1ml 1XPBS for 10 mins at RT with gentle agitation. To mount slides, the PBS was removed and Vectashield® DAPI-containing mounting media (Vector Laboratories) added to pellet. Organoids were mounted using a plastic pasteur pipette to drop onto a slide before a coverslip was applied. Imaging was conducted on a ZEISS Axio Observer 7.1 Microscope and ZEISS LSM880 Airyscan Fast upright confocal microscope.

### Cardiac puncture, plasma extraction, and tissue collection

WT and IFNAR1^−/−^ mice were anesthetized via an i.p injection of Ketamine (100mg/kg) (ilium Ketamil) + Xylazine (10mg/kg) (ilium Xylazil-20). Blood was removed via cardiac puncture and plasma extracted. The colon was removed then the intestine was excised and separated into duodenum, jejunum and ileum with all tissues snap frozen in liquid nitrogen and stored at −80°C.

### RNA isolation and cDNA synthesis

RNA was isolated from intestinal tissues (approximately 50-100mg) using TRIzol (Invitrogen, 15596018). Organoid samples were harvested using Corning® Cell Recovery Solution (Corning®, 354253) and pelleted before using RNAeasy mini kit (Qiagen, 74104) as per manufacturer’s instructions. RNA samples were DNAse treaeted with an Ambion TURBO DNA-free™ Kit (Life Technologies), as per manufacturers guidelines. RNA was reverse transcribed into cDNA using a High-capacity RNA-to-cDNA Reverse Transcription Kit (4368814, Applied Biosystems)

### Real time quantitative polymerase chain reaction (RT-qPCR)

All qPCR was performed in triplicate in standard 384-well plates (4309849, Applied Biosciences, Scoresby, VIC, Australia), with real time quantitative gene expression determined using using Taqman probles (**Table 2**) and analysed by the Quant Studio 6 Flex Real-Time PCR System (Invitrogen).

### Western blot analysis

Western blot analysis was performed on 50μg of protein. Membranes were incubated with the primary antibodies **(Supp. Table 3)** for 24 hours at 4°C and secondary antibodies for 1.5 hours at RT. Signals detected using an ECL Prime Detection kit (GE Healthcare Life Sciences) and visualised with a ChemiDoc™ imaging XRS+ system (Bio-Rad).

### B16 HEK-Blue murine IFN α/β reporter cell line

A B16 HEK-Blue murine IFN α/β reporter cell line (Invivogen) was used as previously described^17^ to determine IFN production in intestinal organoid cultures.

### Enzyme Linked ImmunoSorbent Assay (ELISA)

A Mouse TNF-alpha Duoset ELISA (R&D Systems, DY410) was used to analyse TNFα levels in mouse plasma and in protein samples extracted from WT and IFNAR1^−/−^ mice gut tissues.

### Statistical analysis

All data presented is expressed as the mean ± SEM. All graphical representations and statistical analysis were carried out using GraphPad PRISM® (version 5.03). qPCR data present was analysed using the Comparative CT Method (ΔΔCt)^18^ and is expressed as fold change (compared to vehicle control). B16-Blue assay was analysed using a two-way ANOVA, with Bonferroni post-hoc test. IHC comparisons were analysed via two-way students t-test. For all tests p≤0.05 was accepted as being statistically significant.

## Results

### Expression of phosphorylated α-Syn is elevated in the GIT of aged WT, but not IFNAR1^−/−^ mice at 6-months p.i. of α-Syn PFFs into the striatum

To determine if age-related increases in type-I IFNs influence the brain to gut transmission of α-Syn pathology, young and aged wildtype mice received an intrastriatal injection of α-Syn PFFs or vehicle **(Supp Figure 1)** and their GI tissues were analysed 6 months p.i. Young mice that received an intrastriatal injection of α-Syn PFFs displayed a similar type-I IFN response in the duodenum to that of vehicle controls **(Figure 1)** with no significant differences in the expression of STAT3, IRF3, STAT1, p-Stat1, p-Stat3 or IFN-β. Duodenal expression of GFAP (enteric glial cell marker) and p-αSyn was also unchanged. However, in aged wildtype mice, IRF3 expression was significantly elevated (0.6-fold, **p<0.01) in the duodenum of α-Syn PFF injected mice, as was IFNβ expression (2-fold, *p=0.05), when compared to vehicle **(Figure 2).** There was also an upregulation (2-fold, *p<0.05) in GFAP expression in α-Syn PFF injected mice compared to vehicle control. Western blot analysis of phosphorylated S129 α-Synuclein levels (detecting both human (exogenous) and mouse (endogenous) α-Syn) confirmed a significant upregulation (2-fold, **p<0.01) in the duodenum of α-Syn PFF injected mice. This upregulation was confirmed by immunofluorescence **(Figure 2C)** with p-αsyn puncta present in the villus of the duodenum. Interestingly, this αSyn pathology was also seen in young α-Syn PFF injected mice **(Figure 1C)** that also displayed altered architecture of the intestine. In the young α-Syn PFF injected mice, there was a significant decrease in the villus length accompanied by a significant decrease in crypt length compared to vehicle control animals **(Supp Figure 2).** Young PBS injected mice had a mean villus length of 571µm compared to young α-Syn PFF injected mice (389.7µm, ****p<0.0001). An analysis of crypt length also confirmed a reduction in the average crypt length (60.47µm) in α-Syn Young PFF injected mice compared to PBS injected mice (63.37µm). An analysis of the aged cohort identified no significant differences between PBS and α-Syn PFF injected mice **(Supp Figure 2).**

**Figure 1.**
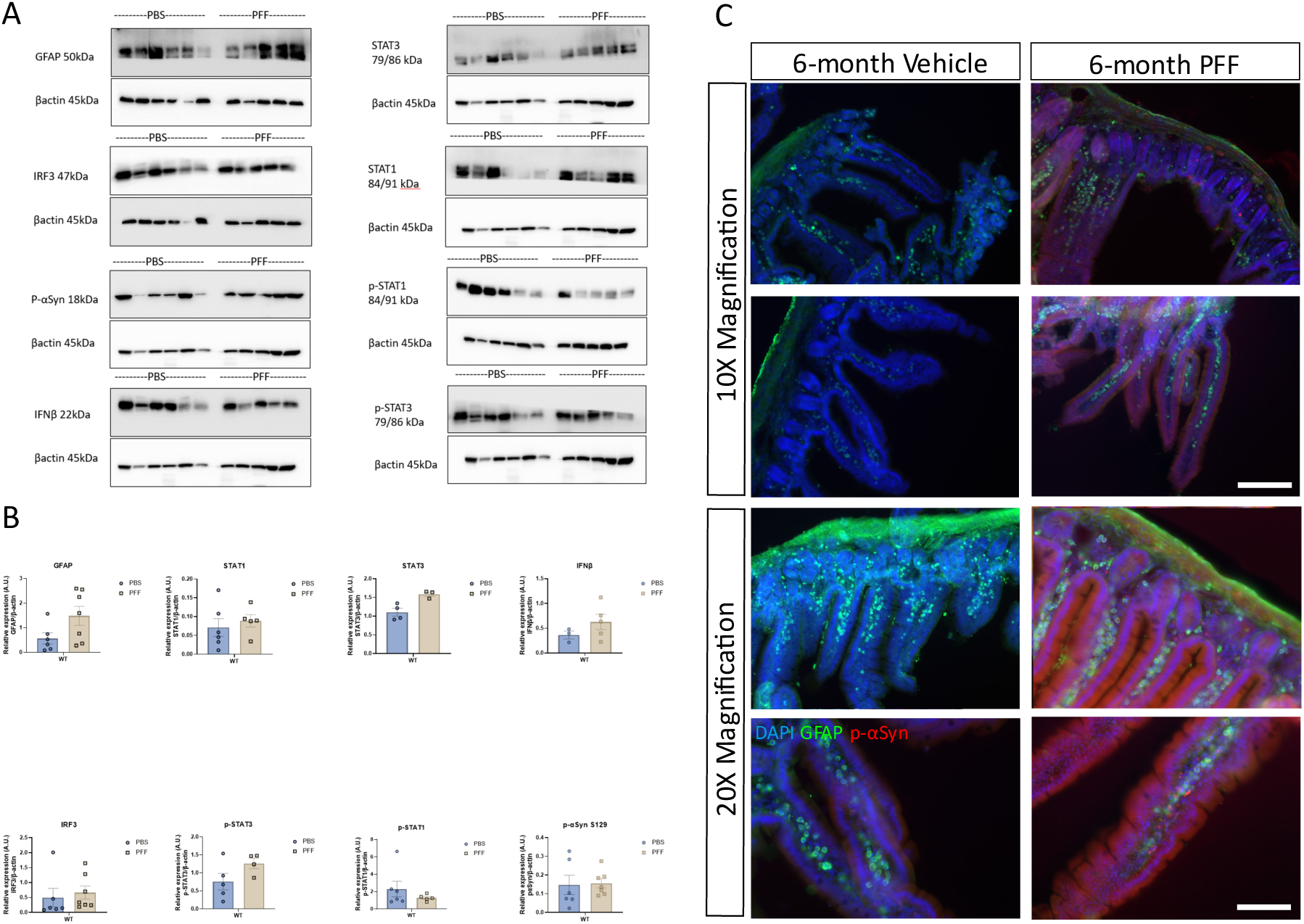
Intrastriatal injection of α-Synuclein PFFs does not induce p-αSynuclein staining or a proinflammatory response in the GI tract of young WT mice at 6-months post injection. Young wildtype mice (10-12 weeks of age at injection) received an intrastriatal injection of α-Syn PFFs (or vehicle), before duodenal tissue was excised at 6-months post injection for analysis by western blot (Panels A, B) or immunohistochemistry (Panel C). A, B; Values expressed relative to β-actin as expressed as mean±SEM, n = 5-6, representative immunoblots of 3 experiments. C; Immunofluorescence images of duodenum from young mice that had received an intrastriatal injection of PBS (vehicle)/a-Syn PFF (PFF) stained for glial fibrillary acidic protein (green), p-αSyn S129 (red), and DAPI (blue). Scale bar depicts 200µm and 100µm, respectively.

**Figure 2.**
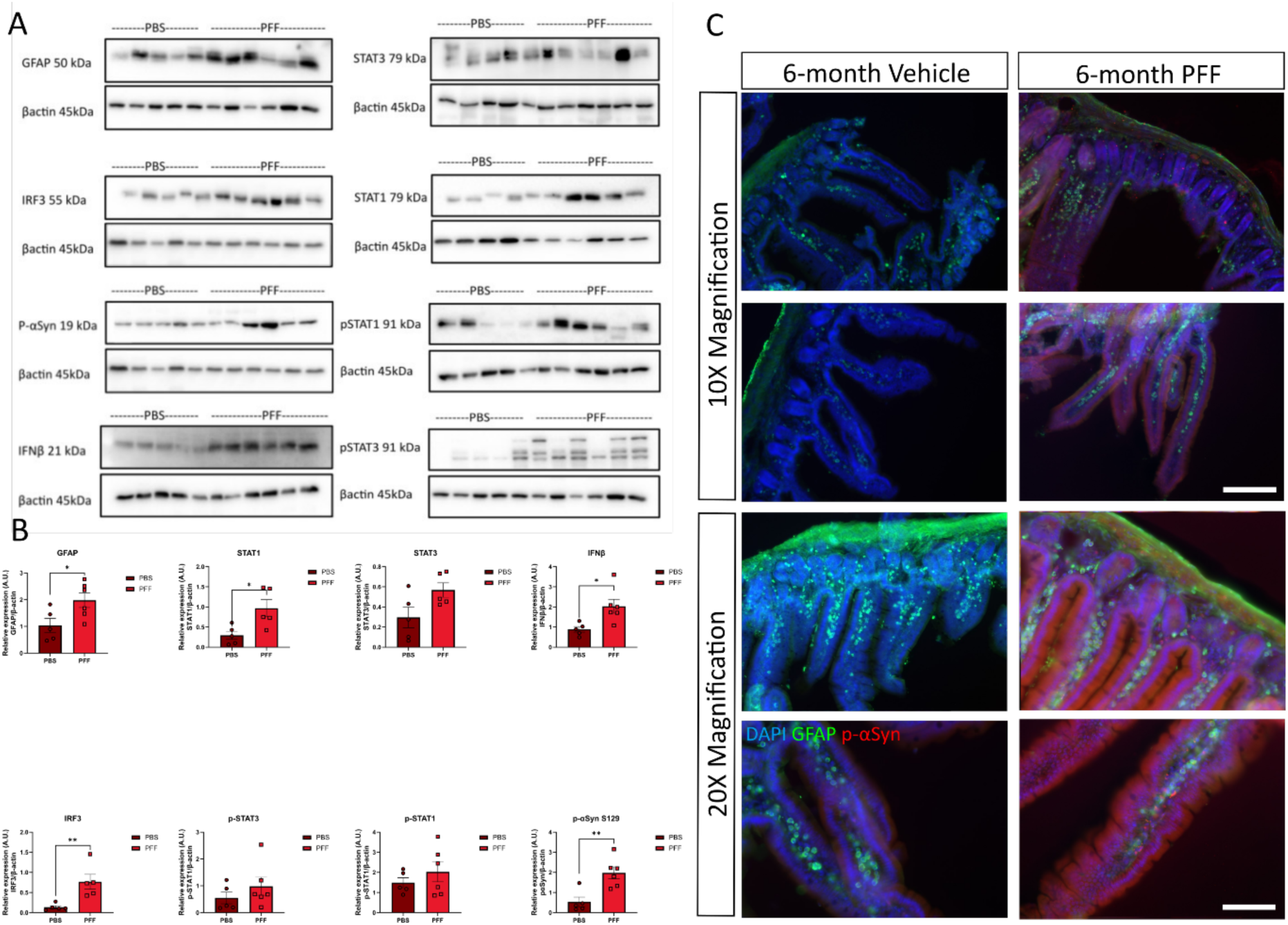
Aging induces p-αSynuclein staining and a proinflammatory response in the GI tract of WT mice at 6-months post intrastriatal injection of αSynuclein PFFs. Duodenum tissue was excised from aged wildtype mice (40-50 weeks of age at injection) 6 months after receiving an intrastriatal injection of αSyn PFFs or vehicle. A, B; Values expressed relative to b-actin and as mean±SEM, n = 5-6, images representative of 3 immunoblots. Unpaired student’s t-test, *p ≤ 0.05, **p ≤ 0.01. C; Immunofluorescence images of 30mM duodenal cryosections from young mice that had received an intrastriatal injection of PBS (vehicle) or a-Syn PFFs (PFF) stained for glial fibrillary acidic protein (green), p-αSyn S129 (red), and DAPI (blue). Scale bar depicts 200µm and 100µm, respectively.

In contrast, there were no significant changes in protein expression of GFAP, IRF3, p-αSyn, and total α-Syn in the duodenum of either the young or old IFNAR1^−/−^ mice that had received an intrastriatal injection of α-Syn PFFs (compared to vehicle controls). Furthermore, through immunofluorescence analysis we observed no discernible changes in GFAP expression, and limited p-αSyn staining in duodenal sections taken from young and aged IFNAR1^−/−^ mice. There were also no significant differences in the villus and crypt length between vehicle and α-Syn PFF injected mice in either the young or aged cohort **(Supp Figure 2D).** These findings suggest that ablation of IFNAR1 influences the transmission of α-Syn from the brain to the gut.

### Intrastriatal injection of α-Syn PFFs induces a type-I IFN mediated proinflammatory response in the GIT of aged mice

Previous studies have confirmed that following an intra-striatal injection of a-Syn PFFs, a pro-inflammatory response is induced in the CNS of mice, however less is known about the gut response in this model. This study saw no significant differences in mRNA expression of IRF3, IRF7, TNFα, IL-1β, TGFβ, and STING between vehicle and α-Syn PFF injected young WT mice **(Figure 3)**. In the colon, α-Syn PFFs induced IRF7 expression (compared to vehicle controls), however the expression of all other genes was unchanged. Conversely, a pro-inflammatory response was detected in duodenum tissue from aged mice at 6-months p.i of α-Syn PFFs, with significant upregulations in IRF3 (3-fold, *p<0.05), IRF7 (1.8-fold, *p<0.05), TNFα (4.5-fold, ***p<0.001), and STING (3.5-fold, p=<0.0001) **(Figure 4)**. In the jejunum, significant upregulations in IRF3 (3.2-fold, ****p<0.0001), IRF7 (3.3-fold, ***p<0.001), and STING (4.6-fold, **p<0.01) was detected in the WT α-Syn PFF injected group compared to vehicle controls. mRNA expression in the ileum tissue was unchanged whilst the colon displayed significant upregulations in both IL-1β (2.2-fold, *p<0.05), and STING (11.4-fold, **p<0.01). In contrast, at 6 months p.i, the expression of IRF3, IRF7, TNFα, IL-1β, TGFβ, and STING were unchanged in the duodenum, jejunum, ileum or colon between α-Syn PFF injected mice and vehicle controls in either the young **(Figure 3)** or aged **(Figure 4)** IFNAR1^−/−^ mice.

**Figure 3.**
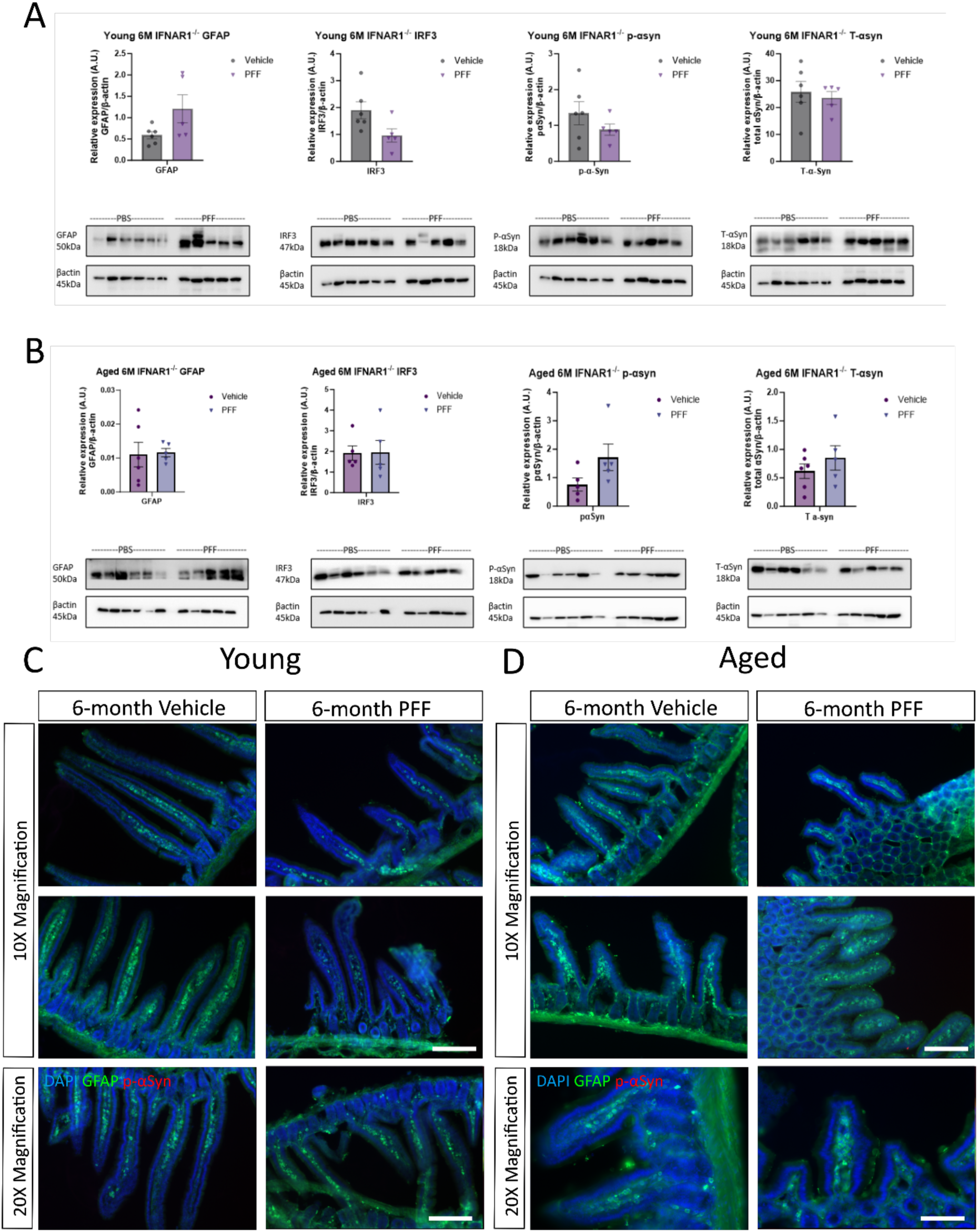
p-αsynuclein staining and proinflammatory response is absent in the GI tract of young and aged IFNAR1^−/−^ mice at 6-months following an intrastriatal injection of αsyn post PFFs. Duodenal tissue was excised from young (10-12 weeks at injection) **(A, C)** and aged (40-50 weeks of age at injection) **(B, D)** IFNAR1^−/−^ mice that had received an intrastriatal injection of αSyn PFFs or vehicle 6-months previously. **A, B**; Western blot analysis values expressed relative to b-actin and as mean±SEM, n = 5-6, images representative of 3 immunoblots. Immunofluorescence images of 30µm duodenal cryosections from young mice (**C**) and aged mice (**D**), stained for glial fibrillary acidic protein (green), p-αSyn S129 (red), and DAPI (blue). Scale bar 100µm and 200µm.

**Figure 4.**
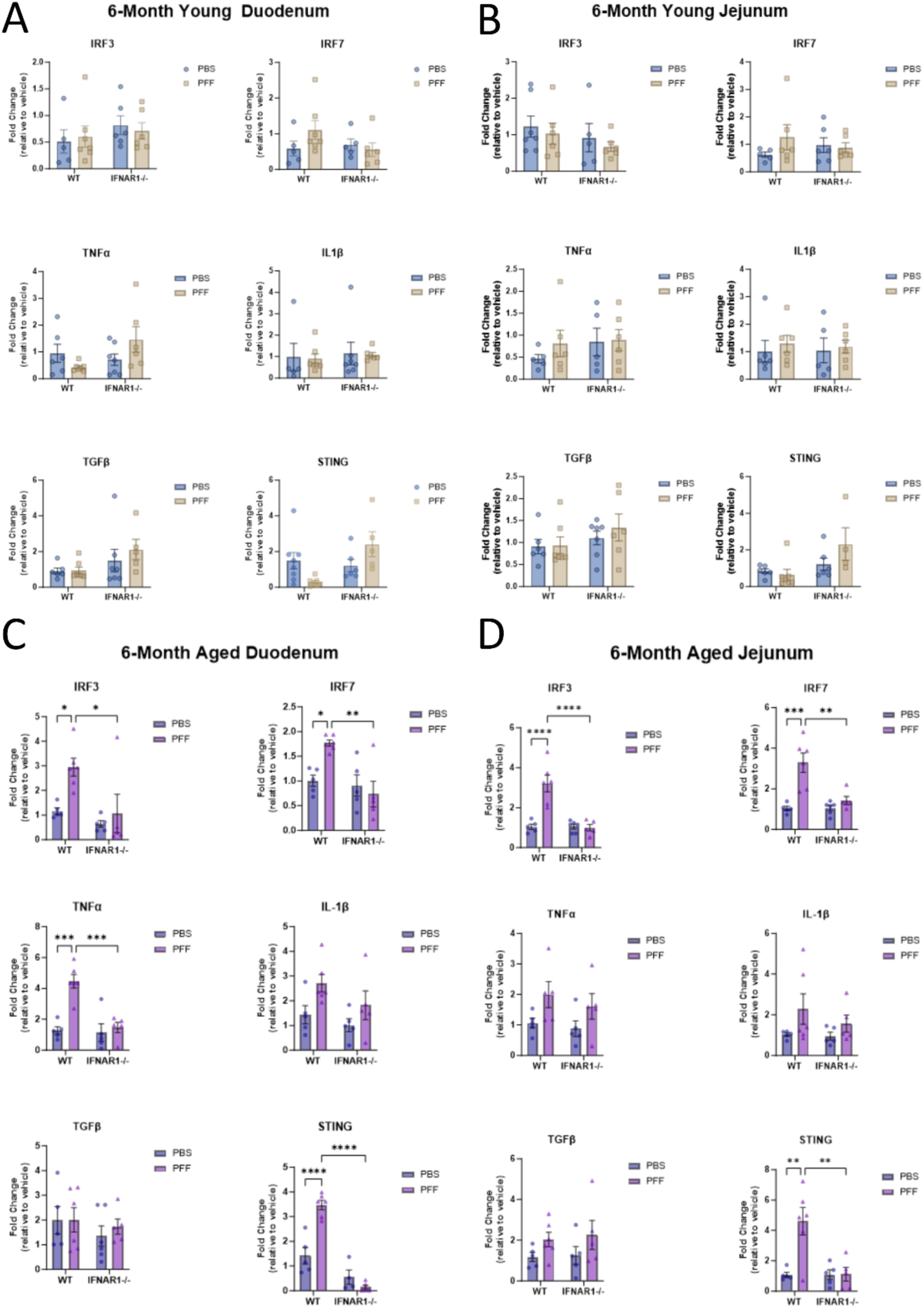
Intrastriatal injection of α-Syn PFF elicits a type-I IFN-mediated proinflammatory response in the duodenum and jejunum of aged wildtype but not IFNAR1^−/−^ mice. mRNA expression analysis of duodenum and jejunum tissue from young (10-12 weeks of age at injection) (**A, B**) and aged (40-50 weeks of age at injection) (**C, D**) WT and IFNAR1^−/−^ mice that had received an intrastriatal injection of α-Syn PFFs or vehicle 6-months previously. Expression levels of Interferon regulatory factor 3 & 7 (IRF3/7), Tumour Necrosis Factor Alpha (TNFα), Interleukin 1-beta (IL-1β), Transforming Growth Factor Beta (TGFβ), and Stimulator of Interferon Genes (STING) were determined as a ratio of beta-2-microglobulin (B2M) expression. Values expressed as fold change (relative to vehicle) and as mean±SEM, n=6-9, two-way ANOVA, Tukey’s Multiple Comparison’s test, *p≤0.05, **p≤0.005, ***p≤0.001, **** p≤0.0001.

### α-Synuclein PFFs are closely localised with enteroendocrine cells within intestinal organoids

The accumulation of α-Syn in neurons and enteric glial cells (EGCs) of the ENS has been reported in GI biopsies from patients with PD with evidence of inflammation also present.^9, 19^ In both mouse and human *in vitro* and *in vivo* models, α-Syn has been shown to colocalise with enteroendocrine cells (EEC).^8^ To further investigate the α-Syn induced proinflammatory response and the contribution of the type-I IFNs, an *in vitro* intestinal organoid model was established. Organoids are a medium throughput screening tool to understand the gastrointestinal epithelium in various disease-specific contexts. In this study, intestinal organoids were used to understand type-I IFNs role in modulating the gastrointestinal epithelial response to α-Syn PFFs.

Murine intestinal organoids were treated with 1μg/ml of human α-Syn PFFs for 48 hours, with immunofluorescence confirming wildtype and IFNAR1^−/−^ organoids had taken up the exogenous a-Syn with total α-Syn staining **(Figure 5A).** Staining with a p-α-Syn S129 (human and mouse specific) antibody **(Figure 5B, C)** was evident within the wall of the intestinal crypt, localising close to the Chromogranin A positive EECs within the intestinal epithelial wall. There did not appear to be any gross discernible differences in the localisation of α-Syn staining between WT and IFNAR1^−/−^ organoids, however western blot analysis did identify reduced levels in the knockout cultures. Relative to vehicle treated organoids, α-Syn PFF-treated wildtype organoids displayed a significant increase in p-α-Syn levels (**p<0.0016) at 48h compared to IFNAR1^−/−^ organoids. This suggests both cultures can take up exogenous α-Syn, however there are differences in its phosphorylation (and potentially aggregation) which may also be true of the endogenous α-Syn species.

**Figure 5.**
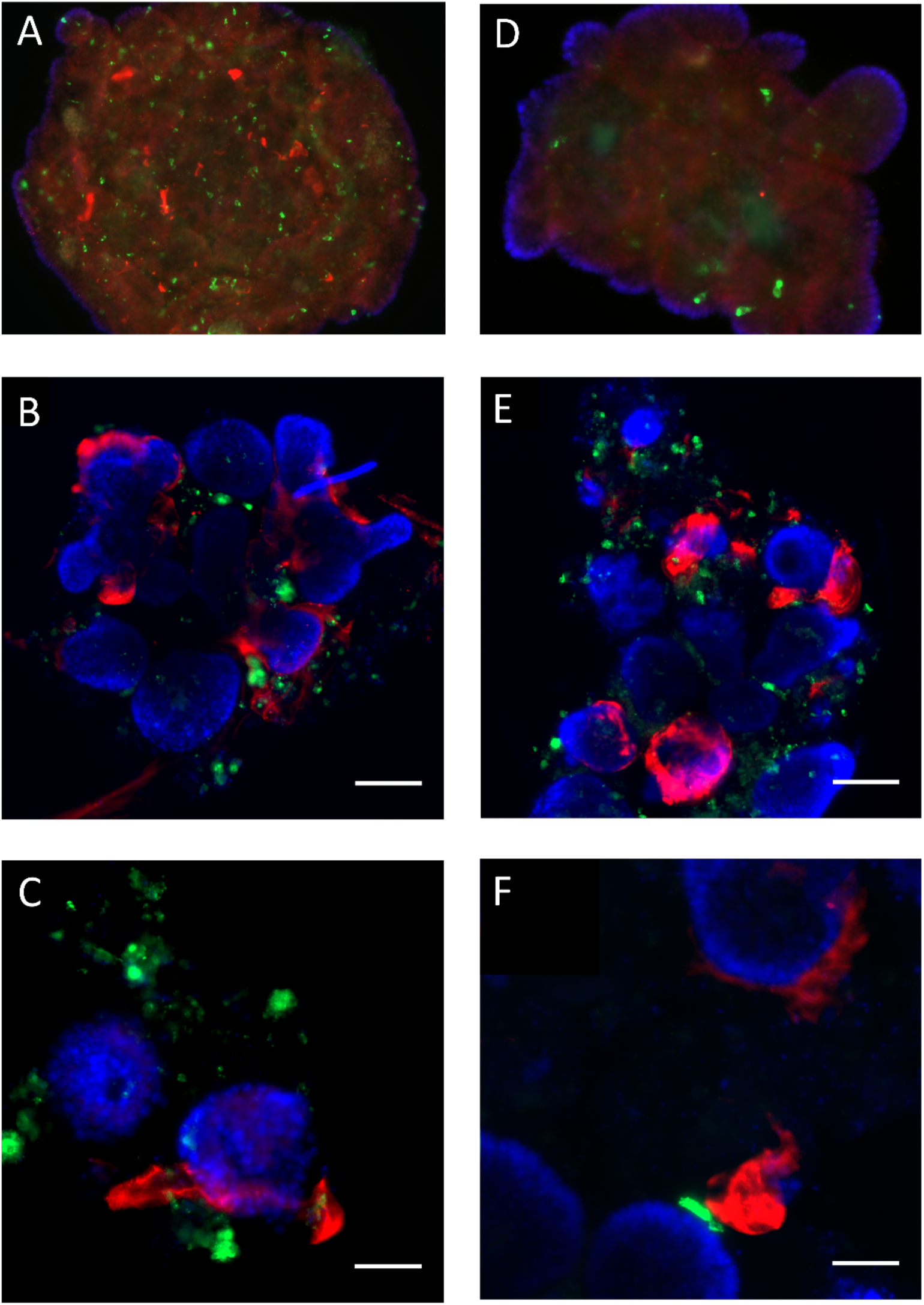
Treatment of wildtype and IFNAR1^−/−^ mouse intestinal organoids with α-Syn PFFs leads to elevated phospho-α-Syn staining near enteroendocrine cells. Representative immunohistochemical images of wildtype (**A, B,** & **C**) and IFNAR1^−/−^ (**D, E,** & **F**) mouse intestinal organoids (passage 1, day 7) treated with 1µg/ml of human α-Syn PFFs for 48-hours. **A**, **D;** Organoids stained with antibodies to the EEC marker, Chromogranin-A (green) and total alpha synuclein (red) and co-stained with DAPI. **B**, **E;** Organoids stained with antibodies to the EEC marker, Chromogranin-A (green) and phospho-Ser129 α-Synuclein (red) and co-stained with DAPI. **C** & **F** Higher power images of **B, F** showing localisation of α-Syn (red) to EEC (green) Scale bar 200µm (A, B, D, E) and 100µm (C, F). Representative images of 3 separate experiments.

**Figure 6.**
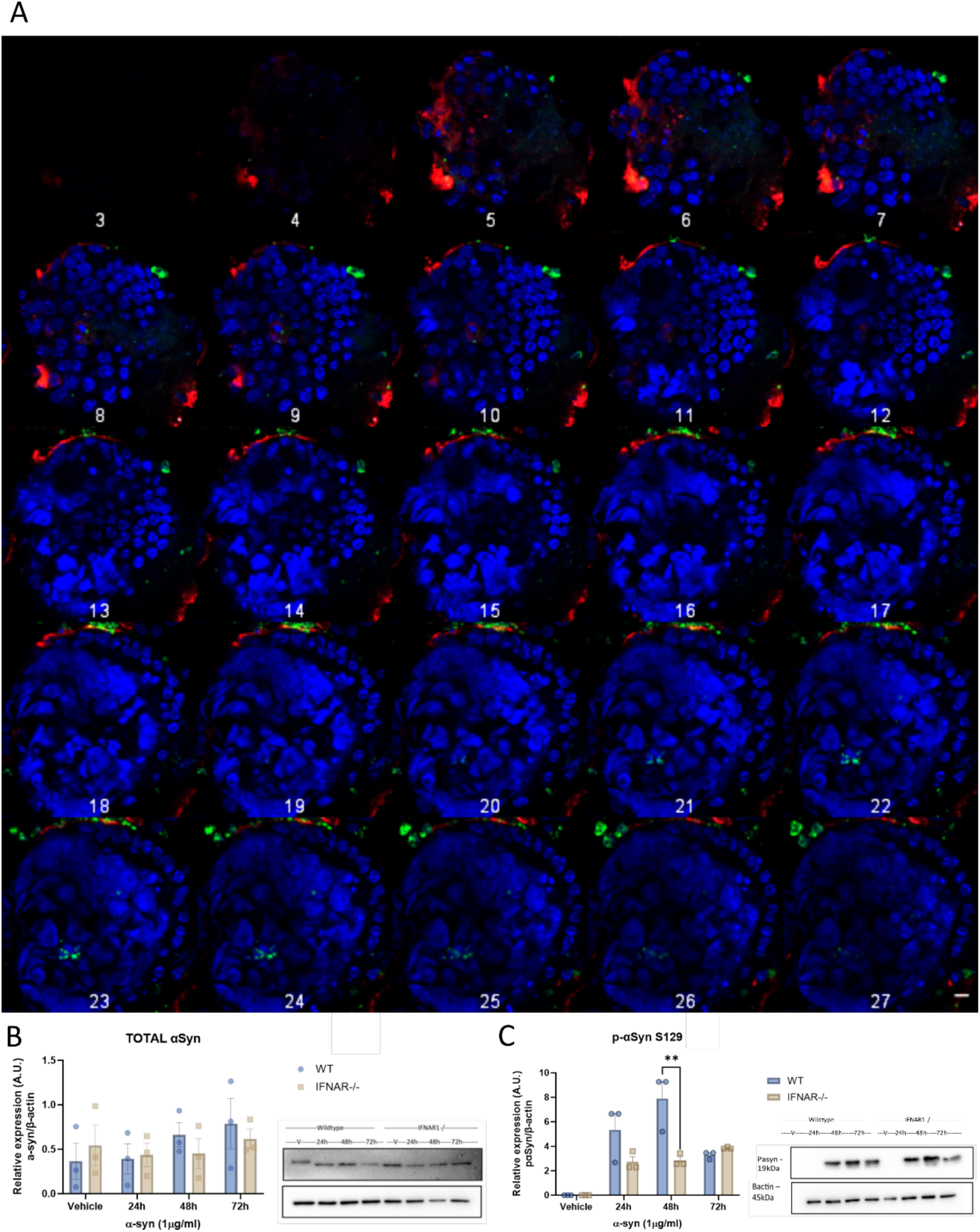
Treatment of wildtype and IFNAR1^−/−^ mouse intestinal organoids with α-Syn PFFs leads to an upregulation in p-S129 α-Synuclein levels. **A,** Wildtype organoids were treated with 1µg/ml α-Syn PFFs for 48h before being stained with anti-α-Synuclein (red), and anti- Chromogranin A (green) and DAPI (blue). Representative images of a Z-stack showing penetration of α-Synuclein into the crypt of the organoid. First slice (1) shows Z position of 0.00nm, each slice afterwards is a Z position of 839.41nm, with a final Z position of 21.82µm. **B, C,** Western blot analysis of wildtype (WT) and IFNAR1^−/−^ organoids treated with 1µg/ml α-Syn PFFs for 24, 48 and 72h. Blots were stained for total alpha synuclein (B) or phosphorylated alpha synuclein (C)) levels with levels expressed as a ratio to b-actin and relative to vehicle control. Two-way ANOVA, data expressed as mean±SEM, n=3, ** p ≤ 0.05.

**Figure 7.**
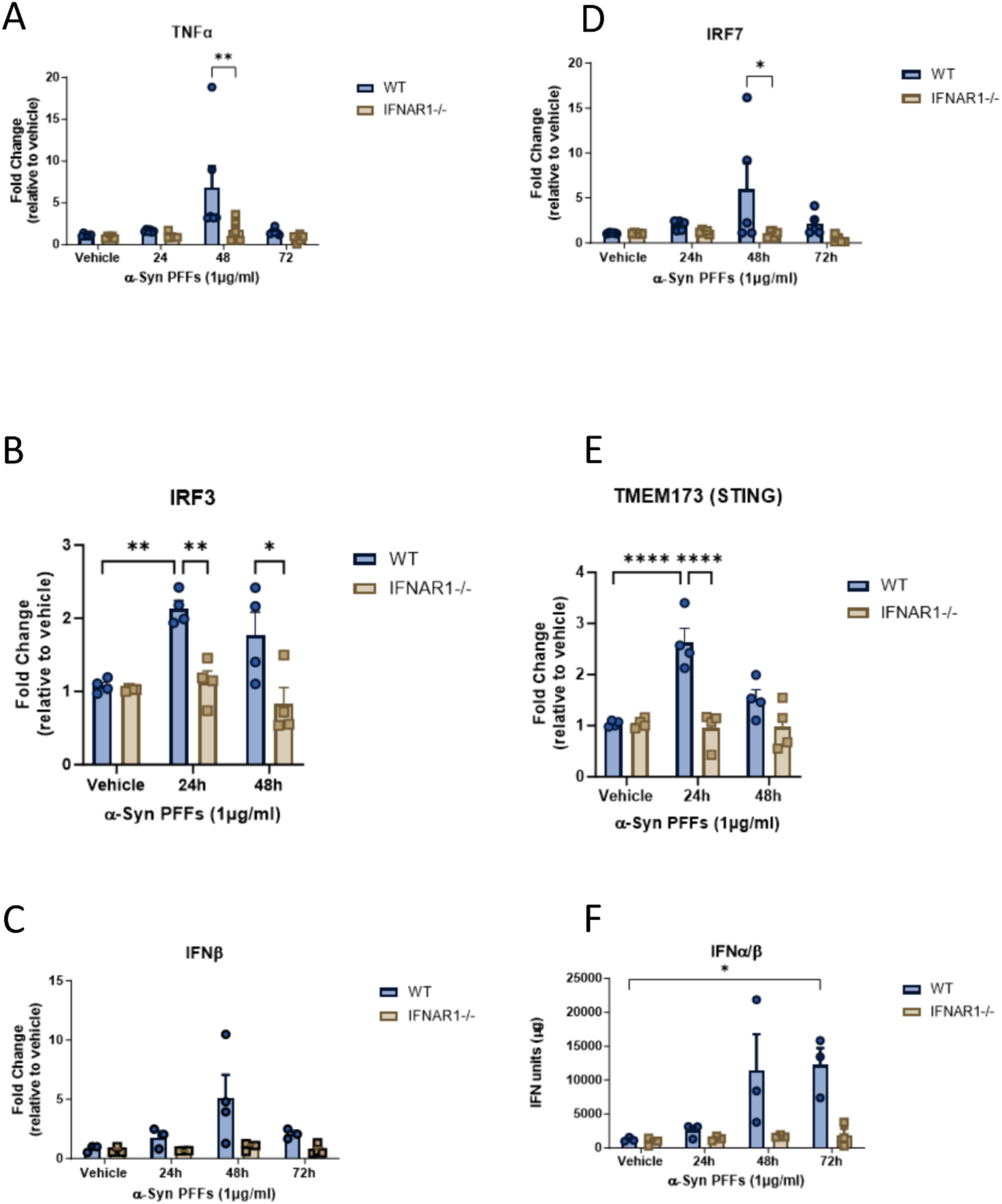
α-Syn PFFs induce a type-I IFN-mediated proinflammatory response in wildtype mouse intestinal organoids that is attenuated in IFNAR1^−/−^ organoids. Wildtype and IFNAR1^−/−^ mouse intestinal organoids were treated with 1µg/ml α-Syn PFFs for 24, 48, and 72h. **A-E**, mRNA expression of Interferon regulatory factor 3 & 7 (IRF3/7), Tumour Necrosis Factor Alpha (TNFα), Interferon Beta (IFNβ), and Stimulator of Interferon Genes (STING). Data expressed as fold change relative to B2M and relative to vehicle control. Data expressed as mean±SEM, Two-way ANOVA, Tukey’s Multiple Comparisons test, * p≤0.05, ** p≤0.01, ****p<0.0001. **F,** A bioactive IFNα/β assay utilising B16 reporter cell lines was used to determine IFN levels in the media from treated organoids. Data expressed in relative IFN units in µg after adjusting for total protein levels. Data expressed as mean±SEM, n=3, Two-way ANOVA, Tukey’s Multiple Comparisons test, *p≤0.05.

### α-Syn PFFs induce a proinflammatory response in mouse intestinal organoids that is type-I IFN dependent

To confirm our *in vivo* findings, and determine if α-Syn PFFs could directly induce a type-I IFN and proinflammatory response in the GIT, wildtype and IFNAR1^−/−^ intestinal organoids were treated with 1μg/ml α-Syn PFFs for 24-72h. qPCR analysis IFNβ expression levels were increased at 48h post α-Syn PFFs treatment in wildtype intestinal organoids (4.811-fold), compared to vehicle controls, with IFNAR1^−/−^ intestinal organoids displaying an attenuated IFNβ response to α-Syn PFFs **(Figure 5).** qPCR analysis also confirmed a significant upregulation in the expression of the downstream mediator, IRF7 in wildtype organoids at 48h (12.684-fold), which was attenuated in the IFNAR1^−/−^ organoids (0.95-fold) **(Figure 5E)**. IRF3 (2-fold, *p<0.05) and STING (2.5-fold, ****p<0.0001) mRNA expression were upregulated at 24h, with both attenuated in the IFNAR1^−/−^ organoids. An upregulation in the type-I IFNs was confirmed using a B16-Blue™ interferon reporter assay. Wildtype organoids displayed robust increases in bioactive IFNα/β expression at 48h (11369.40±5421.14) and 72h (12234.52±2051.19), this response was diminished in the IFNAR1^−/−^ cultures at 48h (1707.81±91.69, *p<0.05) and 72h (1848.14±833.73, *p<0.05) respectively **(Figure 5A-B)**. This attenuated type-I IFN response in the IFNAR1^−/−^ intestinal organoids correlated with a reduced pro-inflammatory response. Wildtype organoids treated with α-Syn PFFs displayed an upregulation in TNFα expression at 48h (compared to vehicle) (5.200±1.882), with this response attenuated in the IFNAR1^−/−^ cultures (1.531±0.589, *p≤0.05) **(Figure 5C-D).**

## Discussion

This study provides evidence for the type-I IFNs in modulating the transmission of α-Synuclein pathology from the CNS to the GIT in a mouse model of PD. This transmission was age dependent with the type-I IFN-mediated inflammatory phenotype only seen in wildtype mice. Type-I IFNs are known to be elevated in aged mice and humans and have previously been linked to aging-related cognitive decline.^13^ Accordingly, we and others have identified the type-I IFNs in both post-mortem human AD and PD brains and mouse models of the diseases.^11, 12, 20^ Although type-I IFN function in the GI tract in homeostatic conditions ^21^ and in injury is well established ^22^, this study is the first to implicate the type-I IFNs in the GIT pathology in PD.

We have previously confirmed an early upregulation in CNS expression of the type-I IFNs in the MPTP mouse model of PD.^11^ IFNAR1^−/−^ mice display reduced neuroinflammation and neuroprotection against nigral cell death and locomotor deficits induced by MPTP. The MPTP model has several limitations; it is an acute model involving the i.p injection of MPTP (4 x 2 hourly doses) and the subsequent analyses at 21 days p.i., therefore it is not a true representation of the human disease progression. The MPTP model also displays an absence of α-Syn pathology within the CNS. The i.p injection of MPTP precludes any specific brain-gut transmission to be identified, therefore for this study we chose to utilise the intrastriatal α-Syn PFF model that induces a significant CNS proinflammatory response, p-αSyn pathology, nigrostriatal pathway loss and behavioural deficits.^23–25^ It should be noted that these studies have predominately focused on the CNS pathology and subsequent motor symptoms that are seen in the human disease. We hypothesised that the type-I IFN-mediated inflammatory response that we reported in the CNS ^11^ would also be a driver of the GIT pathology and non-motor symptoms that occur in PD.

There is supporting evidence for a relationship between α-Syn, inflammation and disease progression along the gut-brain axis in several PD mouse models. Kishimoto et al. (2019) reported chronic intestinal inflammation in α-Syn mutant mice that presented with accelerated brain neuropathology and motor dysfunction.^26^ Furthermore, increased markers of inflammation (TNFα, IL-6, IL-1β, and IL-10) were seen in both the colon and the brain, consistent with the retrograde transneuronal propagation of α-Syn pathology and neuroinflammation from the gut to the brain. These findings were corroborated by Kim et al. (2019) who reported the ability of α-Syn PFFs to induce inflammation and pathology through the transportation of aggregated α-Syn from the gut to the brain.^27^ Other studies have confirmed that inoculation of α-Syn PFFs into the intestinal tract of both mice and rats leads to aggregation of α-syn primarily near the injection site, but also in areas of the brain.^28,29^ In addition, Challis et al (2020) reported a more severe phenotype in aged mice that had received a gut injection of α-Syn PFFs.^30^ Specifically, they found increased p-α-Syn staining in the brainstem and midbrain, compared to young, injected mice. Moreover, the researchers found that aged mice also presented with decreased striatal dopamine, when compared to young, injected mice.

To our knowledge however this is the first time it has been shown that aged mice exhibit greater brain-gut transmission of α-Syn pathology and an exacerbated proinflammatory GI phenotype. Immunohistochemical and western blot analyses identified elevated phosphorylated ser129 α-Syn levels in the duodenum of aged mice **(Figure 2)** that was not seen in young mice **(Figure 1)** at 6-months p.i of α-Syn PFFs into the striatum. This phosphorylated α-Syn antibody detected both human and mouse phosphorylated α-Syn on S129A. Similarly, an elevated type-I IFN and pro-inflammatory response was only seen in the duodenum of aged WT mice. Increases in proinflammatory cytokines (IFNɣ and TNFα) are upregulated in the human PD GIT and have been reported in the GIT of other models of PD including peripheral MPTP injections however this was the first report confirming the type-I IFNs involvement in this response. Type-IFNs are elevated in the aging gut, an in an analysis of mouse intestinal organoid cultures it was specifically shown that the IFN-Stat1 axis drives aging-associated loss of intestinal tissue homeostasis and regeneration.^31^ Furthermore, the group identified an increase in proinflammatory cells in the lamina propria of the aged intestine, that perpetuated the induction of Stat1 activity in intestinal stem cells priming the aberrant differentiation and increase in antigen presentation on the IECs.

The phenotype seen in the aging mouse is indicative of that reported in PD patients, with a worsening of GI barrier function with age.^32^ We postulated that the elevated type-I IFNs in the aging gut were contributing to the exacerbated pro-inflammatory response following an intrastriatal injection of α-Syn PFFs. Indeed, IFNAR1^−/−^ mice displayed an attenuated pro-inflammatory and type-I IFN response with reduced α-Syn staining in the duodenum. Intestinal inflammation has been previously linked to changes in morphology of villus and crypts within the small intestine ^33, 34^, however we only saw these changes in our young mice. In aged mice, no significant differences between treatment groups were identified despite aged mice having lower villus size overall (**Supp Figure 2B)**. However, in the young mice, villus and crypt length in the upper duodenum of α-Syn PFF injected WT mice was reduced compared to their vehicle controls. There was no significant difference in the IFNAR1^−/−^ duodenum **(Supp Figure 2D).** This finding may in part increase our understanding of the type-I IFNs role in the GI epithelium. Type-I IFNs have been linked to intestinal morphology and the composition of the gut microbiota in mice. A study from Brodziak et al. (2013) has shown that the colonic mucosa expression of IFN-related genes was differentially expressed in separate mouse strains after colonisation of commensal microbiota.^35^ Furthermore, an analysis of the microbiota of mice with selective type-I IFN deletion in intestinal epithelial cells depicted changes in the composition of microbiota, Paneth cell abundance, and ability of the epithelium to regenerate.^36^ In addition, Thompson et al. (2010) reported that mice lacking IRF9, which are unable to respond to type-I IFNs, displayed a higher temporal variation of microbiota, accompanied by an increase in the number of T cells and neutrophils in the GIT.^37^ In addition to these changes, it has been presented that in the brain throughout aging there is a type-I IFN signature that is shown negatively affect brain function.^13^ The study reported that blocking IFN-I signalling within the aged brain partially restored cognitive function and hippocampal neurogenesis. In the enteric nervous system, it has been shown that impaired type-I IFN induction in aging impairs viral clearance and promotes immunopathology following infection.^38^ These findings taken together may elucidate a functional decline of both the brain and the gut in aging.

To further determine the effects exogenous α-Syn PFFs have on intestinal morphology, a gut organoid approach was taken. Immunohistochemistry confirmed similar staining patterns of human α-Syn PFFs in both wildtype and IFNAR1^−/−^ murine intestinal organoids, with close localisation of anti-α-Syn staining with Chromogranin A staining of EECs. EECs form the largest endocrine organ in the body and play a key role in the control of GI secretion and motility, regulation of food intake, and glucose homeostasis.^39^ They also have an intrinsic connection to the gut microbiota and have been shown to be modulated in response to short chain fatty acids (SCFAs) and lipopolysaccharides (LPS).^40^ The EECs are complex sensory sentinels of the intestinal environment. They are involved in the detection of inflammation via toll like receptors^41^, and microbiome dysbiosis through microbial metabolites.^42^ In addition, EECs increase crypt cell proliferations via increased growth factor release.^43^ EECs also influence cytokine and antimicrobial production from enterocytes and Paneth cells, whilst maintaining tight junctions of the epithelium.^44, 45^ It has also recently been reported that EECs contain a neuropod that innervates towards the enteric nervous system and glia. This neuropod is hypothesised to enable bidirectional communication between EECs and the enteric nervous system.^40^ Clinical evidence in human patients suffering from IBS has reported that the T-cell mediated inflammation present in GIT is coupled with a 5-fold increase in EEC cell number and changes in gut permeability.^46^ Similarly, increased EECs have also been demonstrated in celiac patients and is associated with inflammation of the intestine.^47^ Recently, the contribution of this secretory cell type to the PD gut disturbances has been proposed. Chandra et al. (2017) identified α-Syn staining within EECs which directly connected to α-Synuclein containing nerves – thus forming a neural circuit between the brain and gut in which toxins and other environmental influences may influence α-Synuclein folding in the EECs.^8^ They suggested that this may provide a possible pathway for misfolded α-Synuclein propagation from the gut epithelium to cause inflammation. Our findings showing α-Syn PFFs inducing a proinflammatory response (increased expression of TNFα) in mouse intestinal organoid cultures support this pathogenic mechanism.

The findings from our *in vitro* cultures corroborate our mouse model studies, identifying the type-I IFNs as key mediators of the α-Syn induced proinflammatory response in the GIT. The proinflammatory response induced by α-Syn was coupled with an increase in both IFNβ and IRF7 levels indicating activation of the IFN signalling pathway. It has been suggested that IRF7 plays a key modulatory role in the homeostatic balance of inflammation within the gut. Recently, Qing and Liu (2023) postulated that IRF7 may play a proinflammatory role in promoting expression of IL-5 and IL-13 induced by BCL11B or directly promoting the expression of inflammatory cytokines and chemokines such as IL-6, TGF-β, CCL5 and CXCL10.^48^ Furthermore, IRF7 KO mice when compared to WT, presented with decreased IL-6 gene expression levels, lower expression of pro-fibrotic factors in fibroblasts, less subcutaneous thickness, and milder inflammation response to bleomycin stimulation.^49^

The type-I IFNs orchestrate a series of intracellular events in immune cells and intestinal epithelial cells (IECs) that lead to the resolution of inflammation and allow the regeneration of the intestinal epithelium and restoration of the gut barrier.^21^ Type-I IFNs are secreted by dendritic cells and phagocytes in response to microbial attack/tissue injury.^50^ Furthermore, type-I IFNs may act on Paneth cells to restrict proliferation and favour their differentiation to establish gut barrier permeability.^21^ However, our findings suggest that in response to α-Syn deposition, the type-I IFNs are instrumental to both the acute and chronic inflammatory response in the GIT. It has been shown that intestinal epithelial cells themselves are capable of producing type-I IFNs, as well as other cytokines and chemokines.^51^ In the DSS model of colitis, mice lacking type-I IFN signalling in IECs display expansion of Paneth cell numbers and hyperproliferation of the intestinal epithelium, when compared to WT littermates.^36^We identified significantly decreased intestinal villus length in our α-Syn PFF-injected IFNAR1^−/−^ mice however, this finding was only seen in young, and not aged mice. It is feasible that the changes in morphology in the guts of aged animals are less responsive to α-Syn, when compared to young mice. This hypothesis is supported by animal studies using the α-Syn PFF model.^30^ It has been postulated that younger adult and aged mice have a differing response due to protein homeostatic function declining in the gut with age.^52^ In addition, the ability to eliminate α-Syn aggregates is expected to be affected to promote gut-brain transmission and sensorimotor deficits.^30^

One of the limitations of this study is that due to the age and accompanying health problems of our aged cohort, the timeline was unable to be extended beyond the 6-months analyses to also confirm α-Syn propagation and pathology in our young cohort. In comparison, in assessing gut-brain transmission, Kim et al. (2019) analysed brains for α-Syn pathology at 10 months post injection.^27^ It should be noted that when validating our model, small but significant motor deficits as assessed by Digigait analysis were identified in the young WT, but not IFNAR1^−/−^ mice at 3 months p.i, (Chen et al., unpublished data). This study chose to focus solely on the GIT in this model and confirmed the type-I IFNs regulate the α-Syn induced neuroinflammatory response implicated in the propagation of pathology along the gut-brain axis. Age-related elevations in type-I IFNs were shown to exacerbate this disease progression, supporting previous studies implicating the type-I IFNs in neuropathologies such as AD.^13^

To our knowledge, this study is the first to use mouse intestinal organoid cultures as a model of GIT dysfunction in PD and allowed us to directly confirm an α-Syn-induced inflammatory response. The use of intestinal organoids in the PD field has not been reported until a recent review paper by Reiner, Sapir, & Parichha, (2021) which discussed future possibilities of using multi-organ systems to model both peripheral and central aspects of PD.^53^ The potential of these systems to facilitate the study of the gut-brain axis in PD *in vitro* and their use for more high-throughput therapeutic approaches to target aspects of disease was discussed. More recently, a paper by Chandra et al. (2023) used intestinal organoids from A53T synuclein mice with isolated nodose ganglion neurons from SNCA^−/−^ mice to understand the transfer of alpha-synuclein between these cell types.^54^ They confirmed human α-Syn transferred between the intestinal organoids and vagal neurons in this co-culture model, with α-Syn pathology moving through vagal neuron processes from cell-to-cell. Our findings specifically demonstrate that α-Syn PFFs are taken up by intestinal organoids, and intestinal organoids mount a type-I IFN-mediated immune response to PFFs Our study is to our knowledge, the first instance it has been shown that intestinal organoids produce an inflammatory response to α-Syn PFF.

Our findings identify a key role for the type-I IFNs within the intestinal epithelium in modulating inflammatory mediators in response to toxins or stressors such as α-Syn. In conclusion, our data confirms that α-Syn pathology in the brain is sufficient to drive an inflammatory response in the GI tract, that is exacerbated by age-related elevations in the type-I IFNs. Future *in vivo* studies will build on our findings from our intestinal organoid cultures to determine if the α-Syn association with the intestinal epithelium and subsequent type-I IFN inflammatory response can be therapeutically targeted to limit the disease progression and non-motor symptoms in PD.

## Acknowledgments

The authors would ike to thank A/Prof Eric Hanssen for assistance with the Transmission Electron Microscopy images and the Biological Optical Microscopy Platform (BOMP) with the Confocal Immunofluorescence microscopy. The authors would also like to thank the Biomedical Science Animal Facility (BSAF) at the University of Melbourne.

## BACKGROUND AND CONTEXT

Type-I interferon (IFN) mediated neuroinflammation is present in the Parkinson’s disease (PD) brain. This study investigated the contribution of the type-I IFNs to the gut response in PD.

## NEW FINDINGS

Type-I IFNs were shown to modulate gut inflammation in response to α-synuclein and exacerbate the progression of α-Syn pathology from the brain to the gut in a mouse model of PD.

## LIMITATIONS

Further studies are required to determine the cellular source within the gut that produces the type-I IFNs in response to α-Syn.

## CLINICAL RESEARCH RELEVANCE

Gut dysfunction is an early clinical symptom in PD patients. This study has implicated a key role for the type-I IFNs in the modulation of the gut-brain axis suggesting that therapeutically targeting this response in the GIT may be a therapeutic target to limit the disease progression in PD.

## BASIC RESEARCH RELEVANCE

This study is the first to implicate age-related increases din type-I IFN signaling in the brain-gut transmission of α-Syn pathology in the mouse. Using gut organoid cultures, the type-I IFNs were shown to be critical mediators of α-Syn-induced proinflammatory conditions, highlighting this model as a powerful in vitro tool to study the gut response in PD.

## Abbreviations

α-syn, alpha-synuclein; CNS, central nervous system; EEC, enteroendocrine cell; EGC, enteric glial cell; ENS, enteric nervous system; GI, gastrointestinal; IFN, interferon; IFNAR, interferon alpha receptor; IL, interleukin; IRF, interferon regulatory factor; PD, Parkinson’s disease; PFF; pre-formed fibrils; SNpc, substantia nigra pars compacta; TNF, tumour necrosis factor; WT, wildtype

## Notes

Conflict of interest statement: The authors disclose no conflicts.

### Competing Interest Statement

The authors have declared no competing interest.

