## Supplemental Tables for "Type-I interferons drive the gastrointestinal inflammatory response in a mouse model of Parkinson’s disease"

### Supplementary Tables

| Cell Type | Antibody | Supplier | Primary dilution | Secondary Antibody |
| --- | --- | --- | --- | --- |
| All | Rabbit Anti-alpha-synuclein | Cell Signalling  (#4179) | 1:200 | Alexa Fluor® 488 Goat anti-mouse (A-11001) |
| All | Rabbit Phospho Anti-alpha-synuclein | Cell Signalling  (#23706) | 1:200 | Alexa Fluor® 594 Goat anti-rabbit (A-21207) |
| Organoids | Purified Mouse Anti-E-cadherin monoclonal | BD Transduction Laboratories™ (#610181) | 1:300 | Alexa Fluor® 594 Goat anti-rabbit (A-21207) |
| Organoids | Rabbit Lysozyme/Muramidase polyclonal | Thermo Scientific (RB-372) | 1:200 | Alexa Fluor® 488 Goat anti-mouse (A-11001) |
| Organoids | Mouse Chromogranin A monoclonal | Santa Cruz (#393941) | 1:250 | Alexa Fluor® 594 Goat anti-rabbit (A-21207) |
| Whole gut IHC | Mouse GFAP monoclonal | Cell Signalling  (#3670) | 1:1000 | Alexa Fluor® 488 Goat anti-mouse (A-11001) |

**Table 1. Antibodies for immunofluorescence analysis**

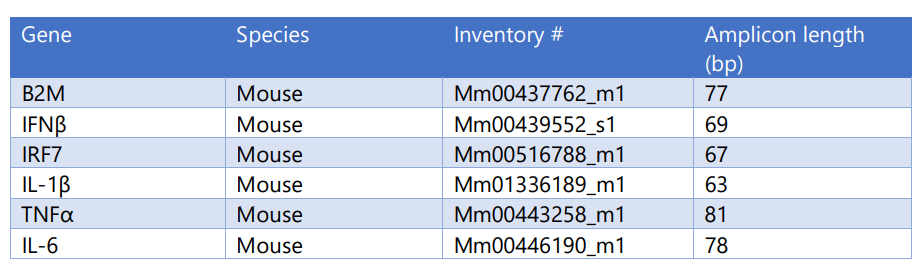
**Table 2. TaqMan probes used for qPCR analysis**

| Antibody | Type | Supplier | Catalogue # | Dilution | Species |
| --- | --- | --- | --- | --- | --- |
| Anti-STAT1 | Primary | Cell Signalling | 9172 | 1:1000 | Rabbit |
| Anti-Phospho-STAT1 | Primary | Cell Signalling | 9167 | 1:1000 | Rabbit |
| Anti-NFκB p65 | Primary | Cell Signalling | 8242 | 1:1000 | Rabbit |
| Anti-Phospho-NFκB p65 | Primary | Cell Signalling | 3031 | 1:1000 | Rabbit |
| Anti-β-Actin | Primary | Sigma-Aldrich | A5441 | 1:1000 | Mouse |
| Anti-GFAP | Primary | Cell Signalling | 3670S | 1:1000 | Rabbit |
| Anti-IRF3 | Primary | Cell Signalling | 4302S | 1:1000 | Rabbit |
| Anti-STAT3 | Primary | Cell Signalling | 4904 | 1:1000 | Rabbit |
| Anti-Phospho-STAT3 | Primary | Cell Signalling | 9145S | 1:1000 | Rabbit |
| Anti-IFNβ | Primary | Santa Cruz | 57201 | 1:500 | Rat |
| Anti-P-S129A αSynuclein | Primary | Cell Signalling | 23706 | 1:1000 | Rabbit |
| Anti-Total αSynuclein | Primary | Cell Signalling | 4179 | 1:1000 | Rabbit |
| Goat Anti-Rabbit Immunoglobulins/HRP | Secondary | Dako | P0448 | 1:1000 | Goat |
| Goat Anti-Mouse Immunoglobulins/HRP | Secondary | Dako | P0447 | 1:1000 | Goat |
| Goat Anti-Rat Immunoglobulins/HRP | Secondary | Abcam | 97057 | 1:1000 | Goat |

**Table 3. Western blot analysis antibodies**
