## Supplemental Figures for "Type-I interferons drive the gastrointestinal inflammatory response in a mouse model of Parkinson’s disease"

### Supplementary figures

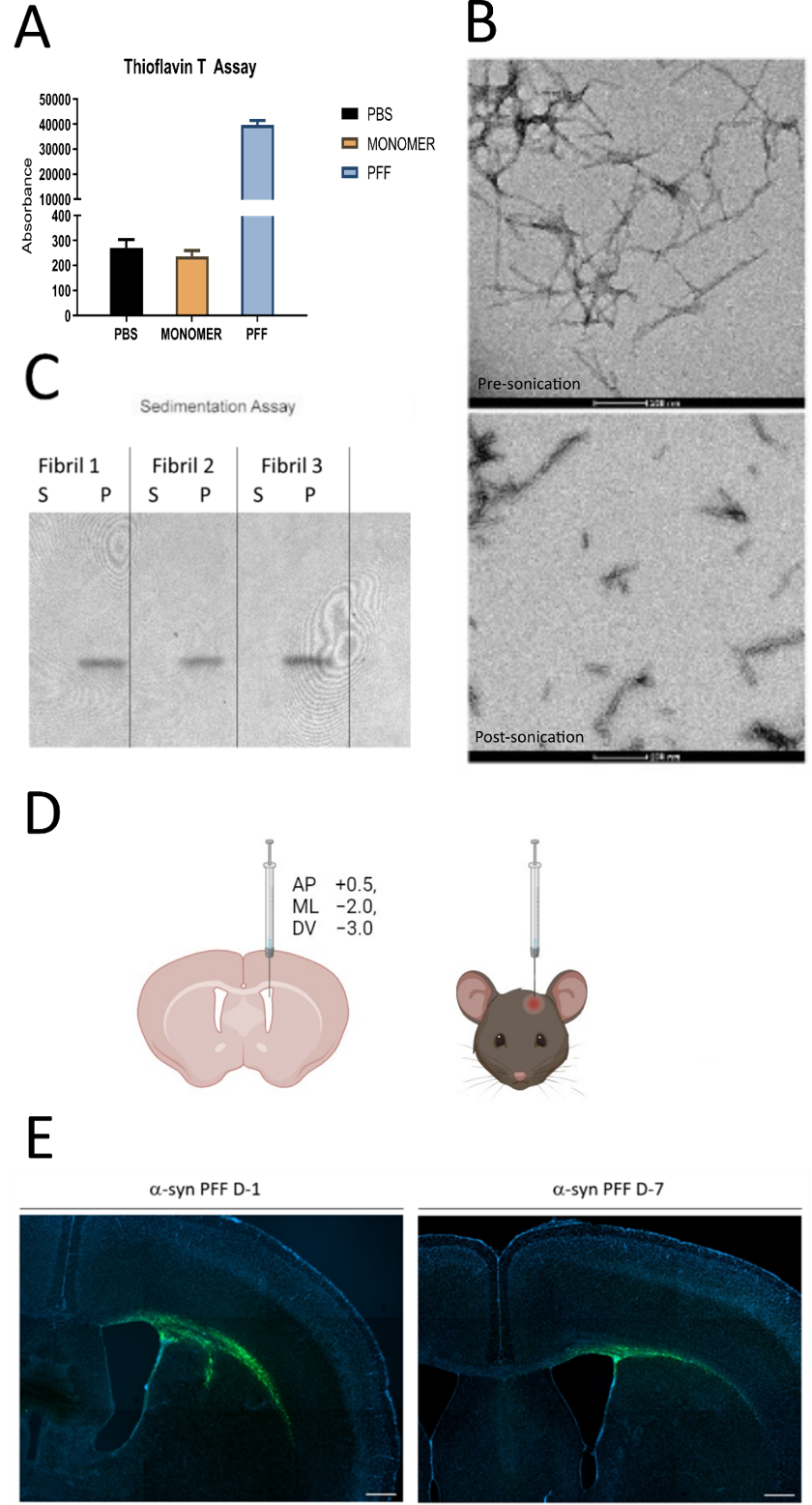

**Supplementary Figure 1. Validation of the alpha synuclein PFF intrastriatal injection model**

**A,** Thioflavin T assay confirming increased β-sheet structures in α-Synuclein PFF, when compared to both monomer and vehicle control. **B,** TEM images of pre- (i) and post- (ii) sonicated preparations of α-Syn PFFs showing decreased length (approximately 50nm) of fibrils. **C,** Sedimentation assay confirming the presence of α-Syn PFFs in the pelleted Representative images of 30µm cryosections of mouse brain at 1- and 7-days post injection with 488-labelled α-Syn PFFs. Co-stained with DAPI. Scale bar =100µm.

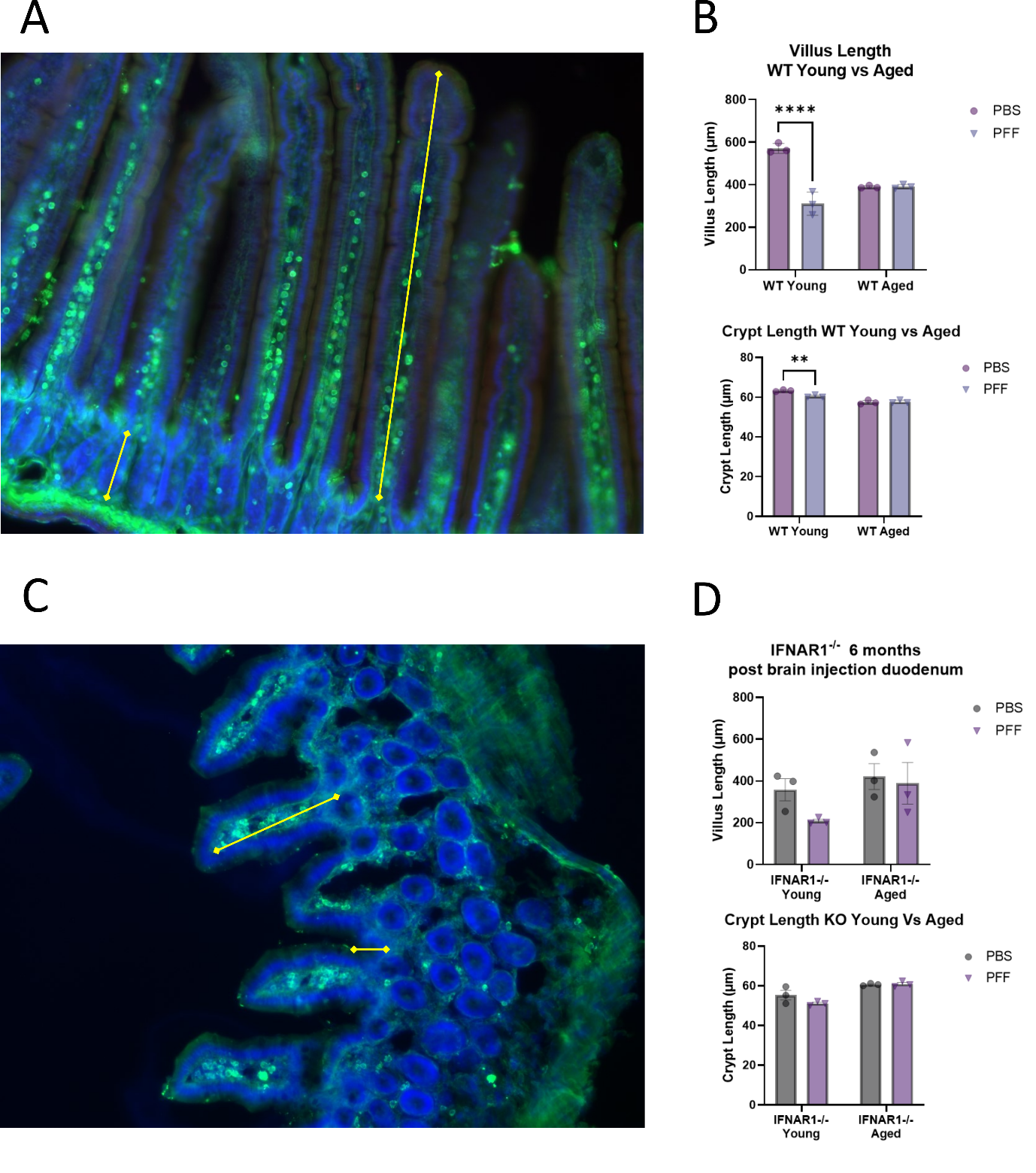

**Supplementary Figure 2. Crypt and villus length is altered in young wildtype, but not IFNAR1^-/-^ mice, following an intrastriatal injection of α-Syn PFFs (6-months p.i).**

Young and aged WT **(A, B)** and IFNAR1^-/-^ **(C, D)** mice duodenal tissue was analysed for villus and crypt length at 6 months post-injection of α-Syn PFFs or vehicle into the striatum. Representative images of 30µm cryosections stained with DAPI (blue) and Anti-GFAP (green) with 3 villus/crypt per cryosection counted, 10 cryosections per mouse, and 3 mice analysed. Data analysed with Two-way ANOVA, Tukey’s Multiple Comparison’s, data expressed as mean±SEM, n=3, **p ≤ 0.01, ****p≤ 0.0001.

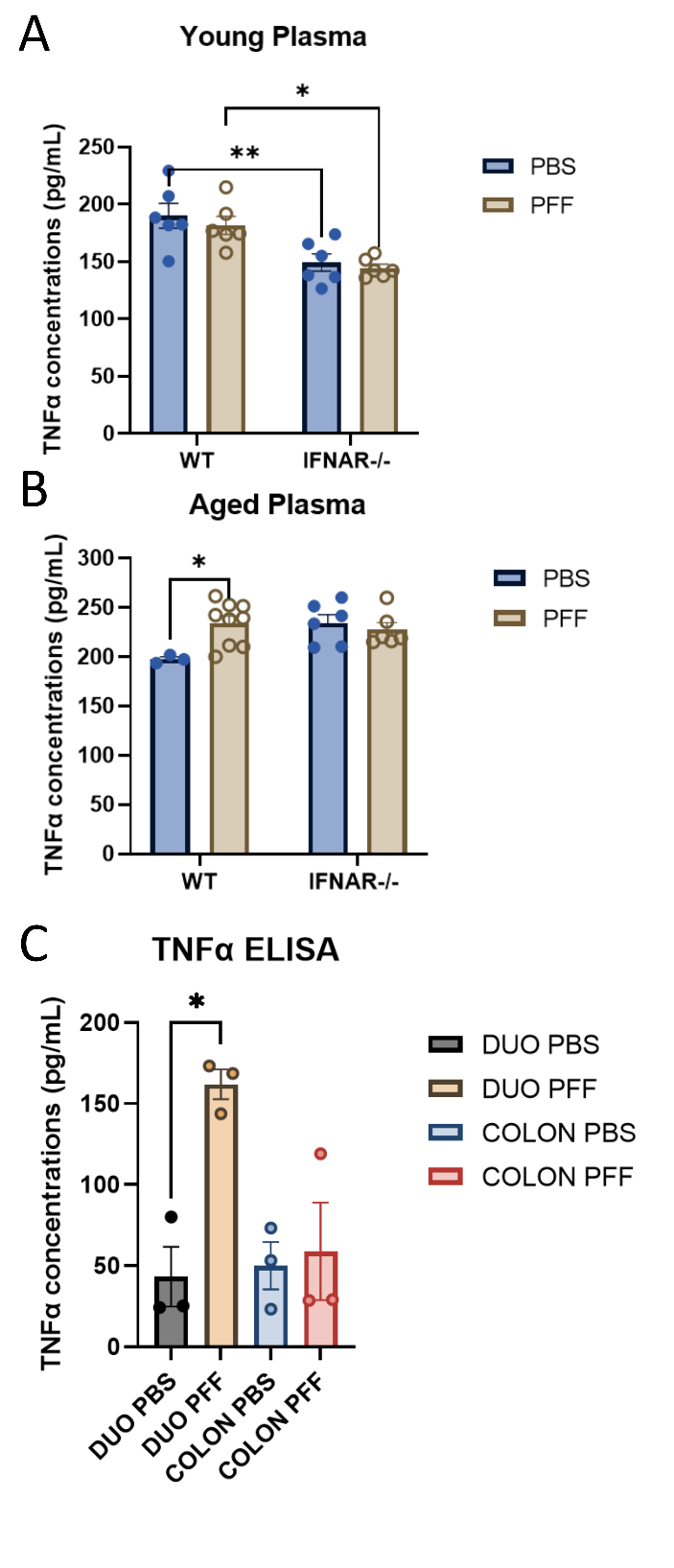

**Supplementary Figure 3. TNFα levels are elevated in the plasma and duodenum of aged wildtype mice, but not IFNAR1^-/-^ mice, following an intrastriatal injection of α-Syn PFFs (6-months p.i.)**

Young (10-12 weeks of age) or Aged (40-50 weeks of age) wildtype and IFNAR1^-/-^ animals were injected with α-Syn PFFs or vehicle and levels of TNFα in plasma (A, B) or duodenum and colon (C) determined by ELISA at 6-months post injection. Data expressed as mean±SEM, two-way ANOVA, Tukey’s multiple comparisons test, * = p≤0.05, ** = p≤0.01.

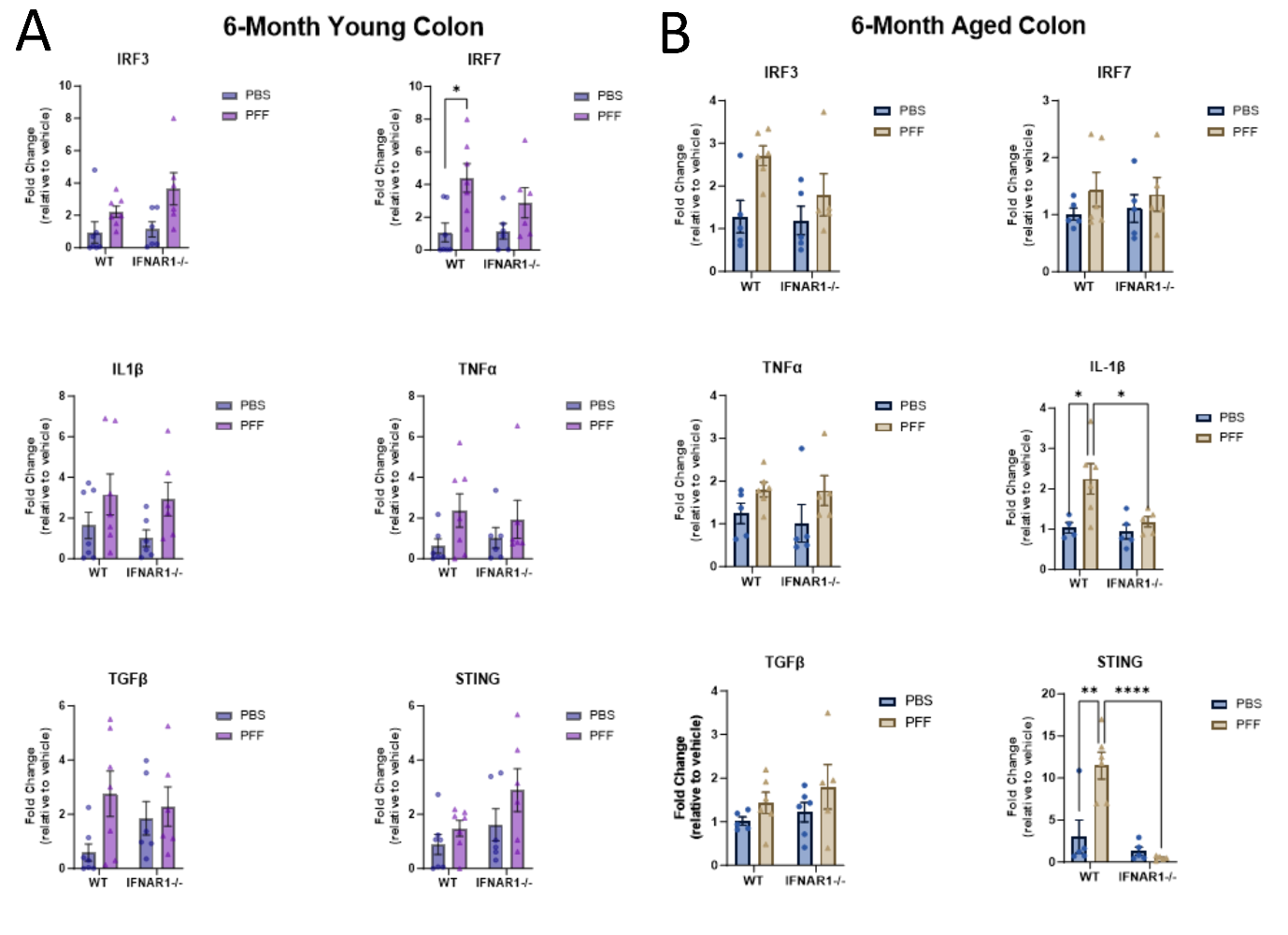

**Supplementary Figure 4. IL1β and STING expression are upregulated in the colon of aged wildtype, but not IFNAR1^-/-^ mice following an intrastriatal injection of α-Syn PFFs (6-months p.i.)**

Colons from young (10-12 weeks of age when injected) **(A)** and aged (40-50 weeks of age when injected) **(B)** WT and IFNAR1^-/-^ mice were analysed at 6-months after receiving an intrastriatal injection of α-Syn PFFs (8µg) or vehicle. mRNA expression of Interferon regulatory factor 3 & 7 (IRF3/7), Tumour Necrosis Factor Alpha (TNFα), Interleukin 1-beta (IL-1β), Transforming Growth Factor Beta (TGFβ), and Stimulator of Interferon Genes (STING) was determined by qPCR analysis. Data expressed relative to the housekeeping gene B2M and as fold change relative to vehicle control, mean±SEM, n=6-10, two-way ANOVA, Sidak’s multiple comparison’s test, *p≤0.05, **p≤0.01, ****p≤0.0001.
